# Impact of AIM2 on HNSCC Development

**DOI:** 10.1101/2024.09.27.615454

**Authors:** Dakota M. Reinartz, Vicente Escamilla-River, Stephanie L. Tribble, Carlos Caulin, Justin E. Wilson

## Abstract

Head and neck squamous cell carcinoma (HNSCC) constitutes 90% of head and neck cancers. HNSCC development is linked to chronic inflammation, while established HNSCC tumors are often immune suppressive. However, both occur through mechanisms that are not fully understood. The cytosolic double-stranded DNA sensor Absent in Melanoma 2 (AIM2) is an inflammasome forming protein that also has inflammasome-distinct roles in restricting tumorigenesis by limited PI3K signaling. Here, we used an experimental mouse model of HNSCC, involving treatment of wild type (WT) and *Aim2^−/−^*mice with the carcinogen 4NQO in drinking water. Compared to WT mice, 4NQO-treated *Aim2^−/−^* mice exhibited larger tumor sizes and increased tissue dysplasia. 4NQO-treated wild type and *Aim2^−/−^* mice displayed similar tongue *Il6, Tnf, Il1b, Il12, and Il10* expression and no consistent differences in PI3K or inflammasome activation, suggesting AIM2 may not regulate these factors during HNSCC. Instead, *Ifng* and *Irf1* was elevated in 4NQO-treated *Aim2^−/−^* mice, suggesting AIM2 restricts IFNγ. In line with this, RNA-sequencing of total tongue RNA from 4NQO-treated mice revealed *Aim2^−/−^* mice had enhanced expression of genes related to the MHC protein complex, cell killing, and T cell activation compared to wild type mice. In addition, we observed increased macrophage infiltration into the tongue epithelium of 4NQO-treated *Aim2^−/−^* mice. Lastly, using *Aim2^−/−^*/*Rag1^−/−^*-double deficient animals, we found that the adaptive immune compartment was necessary for the enhanced tumorigenesis during AIM2 deficiency. Taken together, these findings suggest AIM2 limits the progression of oral tumor development partially through regulating IFNγ and adaptive immune responses.

## Introduction

Head and neck squamous cell carcinomas (HNSCC) account for 90% of all HNC cases^1^. It is the seventh most common cancer worldwide and accounts for about 325,000 deaths annually^1^. The five-year survival rates for HNSCC is at about 50% for all cases, thus highlighting the need for a better understanding of HNSCC development and progression^2^. The known primary risk factors that lead to HNSCC development are tobacco combined with heavy alcohol consumption^3^ and human papilloma virus (HPV) infection^4^. Genetic mutations in members of the PI3K/Akt pathway the JAK/STAT pathways, and MAPK pathways are among the most frequently mutated signaling pathways in HPV^+^ HNSCC but is also observed in HPV^−^ HNSCC but at a lower frequency^6^. In addition to regulating cellular survival and proliferation, these pathways are also primary signal mediators of inflammatory signaling. Similar to many cancers, increasing evidence also points towards chronic inflammation as a key player during HNSCC tumorigenesis^7,8^. However, the cellular and molecular mechanisms that regulate inflammation during HNSCC are not fully understood.

Inflammation is largely initiated through the activation of pattern recognition receptors (PRRs). These receptors recognize specific molecular patterns frequently found in microbes (known as pathogen-associated molecular patters or PAMPs) and molecules that are released by damaged or dying cells (known as damage-associated molecular patters or DAMPs) ^9^. Upon recognition of these PAMS or DAMPs, these receptors trigger immune responses that initiate localized. There are many different types of PRRs with the most well described being Toll-like receptors (TLRs) and nucleotide oligomerization domain (NOD)-like receptors. Although vital to combating infection, the aberrant or chronic activation of PRRs can contribute to autoinflammatory disorders and many cancers. For instance, elevated TLR4 positively correlates with HNSCC tumor grade. Activation of TLR4 with lipopolysaccharide leads to enhanced proliferation of tumor cells and production of inflammatory cytokines, including Interleukin-6 and IL-8, along with vascular endothelial growth factor and granulocyte macrophage colony-stimulating factor, which support the progression of HNSCC^10^. Other TLRs, like TLR2 and TLR9, are also highly expressed in early-stage HNSCC and are putative predictors of invasive tumor growth^11^. Animals deficient in the intracellular PRR NLRP3 are resistant to experimental HNSCC, indicating that NLRP3 promotes HNSCC tumorigenesis^12,13^. Mechanistically, activation of NLRP3 leads to formation of the inflammasome and production of IL-1β, which promotes early and late stage HNSCC through increasing proliferation of dysplastic oral cells, stimulating the production of oncogenic cytokines and making tumors more aggressive^14^. These findings indicate a crucial role for PRR dysregulation and chronic inflammation in promoting HNSCC, however the contribution of additional PRRs during this process has yet to be explored.

Absent in Melanoma 2 (AIM2) is a PRR that displays important functions during both inflammatory responses and tumorigenesis. AIM2 consists of a DNA-binding HIN200 domain and a pyrin domain, which complexes with the common inflammasome adaptor protein ASC to form an inflammasome^15^. Similar to the NLRP3 inflammasome, this leads to the release of the proinflammatory cytokines IL-1β and IL-18 and an inflammatory form of cell death called pyroptosis. AIM2 also performs inflammasome-independent functions that limit experimental models of inflammation-associated colon cancer via inhibition of PI3K/Akt and suppression of cellular survival, growth and proliferation^16^. In the context of HNSCC, AIM2 expression was found to be upregulated in established tumors, and its expression is correlated with tumor cell radio resistance, metastasis and PD-L1 expression^17^. However, it is unclear how AIM2 impacts tumor development in vivo. While AIM2 regulates multiple factors associated with HNSCC development including IL-1β expression and PI3K/Akt, the contribution of AIM2 during these early stages is unknown.

In this study, we assessed the role of AIM2 during HNSCC development by treating wild type (WT) and *Aim2^−/−^* mice with the tobacco-surrogate 4NQO. 4NQO is a water-soluble carcinogen that induces DNA adduct formation, leading to mutations and histological abnormalities that resemble oral cancer progression in humans^18^. We observed that 4NQO-treated *Aim2^−/−^* mice grew larger oral tumors and displayed greater dysplasia development on their tongue. The increased tumorigenesis observed in 4NQO-treated *Aim2^−/−^* mice was not associated with loss of inflammasome activity or enhanced PI3K/AKT. Instead, RNA sequencing revealed that *Aim2^−/−^* mice displayed heighted IFN-γ transcriptional signature as well as an increase in macrophage infiltration in their tongue tissue. Furthermore, we found that in RAG1^−/−^/*Aim2^−/−^* mice, which lack AIM2 and an adaptive immune compartment, did not recapitulate the elevated tumorigenesis observed in *Aim2^−/−^* mice after 4NQO treatment. However, we detected no measurable differences in tumor burdens in RAG1KO mice reconstituted with WT or *Aim2^−/−^* CD4 T cells or CD8 T cells, indicating lymphocytes are necessary, but not sufficient for AIM2-mediated protection against HNSCC. Altogether, our results show that AIM2 restricts HNSCC development and IFN-γ-associated transcriptional signatures in oral tissue.

## Methods

### Mice

All animal protocols were approved by the University of Arizona Institutional Animal Care and Use Committee (IACUC) in accordance with the US National Institutes of Health Guide for Care and Use of Laboratory Animals. Specific pathogen free (SPF) C57BL/6 mice (wild type; WT) were originally purchased from Jackson Laboratories and bred at the University of Arizona to avoid facility-associated microbiome differences between experimental groups. Mice deficient in AIM2 (*Aim2^−/−^*) on the C57BL/6 background were generated by Ingenious Targeting Laboratory (Ronkonkoma, NY) as previously described^16^ before being established at the University of Arizona. Mice deficient in interferonγ receptor 1 (*Ifngr1^−/−^*) on the C57BL/6 background and RAG1 (*Rag1^−/−^*) deficient mice on the C57BL/6 background were also purchased from Jackson Laboratories. Mice deficient in AIM2 and RAG1 were bred together to generate *Aim2^−/−^* /*Rag1^−/−^* double knockout mice.

All mice were bred under SPF conditions in the University of Arizona Health Sciences animal facility with NIH standard food and water provided ad libitum.

### Experimental Head and Neck Carcinogenesis

For induction of head and neck cancer, mice were given 100 μg/mL 4-Nitroquinoline-1-oxide (4NQO) dissolved in their drinking water for 20 weeks{Vitale-Cross, 2009 #19}. Animal weights were tracked throughout the experiment. At the end of 20 weeks mice were euthanized or when moribund for macroscopic tumor counts and size measurements using calipers. Macroscopic tumor loads were calculated by adding the total sizes of each tumor for each mouse.

For the adoptive T cell transfer studies, total CD4^+^ T cells or total CD8a^+^ T cells were isolated from WT or *Aim2^−/−^*mice using a magnetic cellular isolation kit (CD4^+^, Miltenyi Biotec, 130-104-454; CD8a^+^, Miltenyi Biotec, 130-104-075) and MACS separation columns (Miltenyi Biotec, 103-042-401). A total of 1 x 10^6^ cells were intravenously injected retroorbitally into *Rag1^−/−^* recipients and allowed to reconstitute for 4 weeks before verifying for reconstitution in peripheral blood samples by FACS. Then mice were given 4NQO as described above.

### Histology

For the assessment of tissue pathology, animal tongues were removed and fixed in 4% paraformaldehyde. Tongues were then paraffin embedded, sectioned and placed onto slides by the University of Arizona Tissue Acquisition and Cellular/Molecular Analysis Shared Resource. Tissue was then stained with hematoxylin and eosin and assessed for tissue pathology by a trained investigator blinded to the experimental conditions of the study.

### Immunohistochemistry

Tissue samples were deparaffinized in xylenes and rehydrated in ethanol-water series. Antigen retrieval was performed in 10 mM sodium citrate buffer (pH 6.0). Then, the sections were blocked and incubated overnight at 4°C with the following unconjugated antibodies diluted in 2.5% normal horse serum: anti CD3e, (Invitrogen # CMA1-90582) at 1:100 dilution or F4/80 (Cell Signaling Technology #70076) at 1:500 dilution. The sections were then incubated with anti-rabbit IgG conjugated with HRP (Vector Laboratories # MP-7801-15). The immunoreactions were visualized with a DAB substrate kit (Vector Laboratories # SK-4105). The sections were mounted and imaged. The number of positive cells to CD3 or F4/80 was analyzed using the positive detection script in QuPath (v0.5.0). Manual annotations were made to distinguish the mucosa and submucosa from the muscular regions of the tongue. Before analysis, color deconvolution was applied, and the same vectors were used across all samples. The density of positive cells was calculated within the annotated areas and reported as cells per square millimeter (cells/mm²).

### Protein Expression

Protein lysates were prepared from tongue sections by homogenizing tongue tissue in radioimmunoprecipitation assay buffer (RIPA; Boston Bioproducts, BP-115) containing 1X Complete Protease Inhibitor Cocktail (Roche, 11697498001), 5 mM Sodium fluoride (Sigma-Aldrich, 201154), 1 mM Sodium orthovanadate (Sigma-Aldrich, 450243), and 1 mM Phenylmethanesulfonyl fluoride (Sigma-Aldrich, 78830). Lysates were transferred into pre-chilled 1.5 ml tubes and centrifuged at 21,130 rcf for 10 minutes at 4°C in a microcentrifuge to pellet the insoluble material. The supernatant was then transferred to a new pre-chilled 1.5 ml tube and the protein concentration was determined. The protein concentration of the supernatant from the tongue tissue were measured by Pierce bicinchoninic acid (BCA) assay (Thermo Scientific, 23225). Lysates containing 20-30 μg of protein were prepared with 4x SDS loading dye (Novex, NP0008) containing 20 mg/ml Dithiothreitol reducing agent (Sigma, D9163-5G). Samples were then boiled for 5 min at 98°C and subjected to 10% SDS-PAGE for protein separation. Separated proteins were wet transferred to nitrocellulose membranes by electroblotting for 75 minutes at 100V at 4°C. Blotted membranes were blocked using 5% non-fat milk for 1 hour incubated with primary antibody of interest overnight at 4°C with constant rocking, washed in 1X TBS-T five times, incubated with the appropriate HRP-conjugated secondary antibody for 2 hours at room temperature, followed by an additional five washes with TBS-T. Antigen-antibody complexes were detected with a chemiluminescence reagent kit using either SuperSignal West Pico PLUS (Thermo Scientific, 34578) or SuperSignal West Femto (Thermo Scientific, 34096) and exposure to autoradiographic film. Film was processed using an automatic film developer (Konica SRX-101A). Densitometry was performed to analyze the western blots utilizing ImageJ.

### RNA Isolation and Analysis

Total RNA was isolated from snap-frozen tongue sections utilizing the RNeasy Plus Mini Kit (Qiagen,74136). Briefly, tongue sections were homogenized in RLT lysis buffer containing β-mercaptoetheanol (Sigma, SHBL6769) using a bead beater homogenizer. Homogenates were then passed through the Qiashredder columns and a genomic DNA eliminator column. We then proceeded with RNA isolation according to the manufacturer’s protocol. RNA concentrations were determined using a spectrophotometer. cDNA was synthesized at a 200 μg/μl template using the iScript Reverse Transcription Supermix for RT-QPCR (Bio-Rad, 1706840) and diluted 1:10 with molecular grade water before performing TaqMan RT-qPCR.

### RNA-Sequencing (RNA-seq)

For RNA-seq analysis Total RNA was extracted as described above and was sequenced and analyzed by Novogene (Sacramento, CA).

### Quantitative Real-Time PCR

qPCR analysis was performed on cDNA from tongue section samples using the TaqMan expression assays (Applied Biosystems). The primer targets used in this study include the following: *Tnfa*, *Il1b*, *Il6*, *Il12*, *Il10*, *Ifng*, *Irf1*, *H2Eb1*, *H2Ab1*. Reactions were run in technical duplicates on a Quantstudio 3 System in a 96-well 0.2 ml block. Relative expression was calculated with the ΔΔCT method using *Gapdh* expression as the endogenous control. Relative expression was plotted as the mean +/− SEM.

### In vitro CD4 and CD8 T cell Isolation and Stimulations

Animal spleens were removed and homogenized with frosted microscope slides in RPMI media. Red blood cells were then lysed with ACK lysis buffer (Gibco, A10492-01). Naïve CD4^+^ T cells and naïve CD8a^+^ T cells were enriched using either a naïve CD4 isolation kit (Miltenyi Biotec, 130-104-453) or naïve CD8a isolation kit (Miltenyi Biotec, 130-096-543) and MACS separation columns (Miltenyi Biotec, 103-042-401). Then, 1 x 10^6^ cells were stimulated in an αCD3 antibody (Invitrogen, 16-0031-coated plate at 2 μg/mL with 0.5μg/mL αCD28 antibody (Invitrogen, 16-0281-82), 5 ng/mL murine IL-2 (Peprotech, 212-12), 10 ng/mL recombinant mouse IL-12 (R&D Systems, 419-ML-010/CF) and 1 μg/mL αIL-4 (Invitrogen, 14-7041-85) for 3 days for CD4 cells. For CD8 cells we performed the same stimulus without the αIL-4. After 3 days supernatants and total RNA were harvested to run ELISA and RT-qPCR.

### ELISA

Supernatants from stimulated naïve CD4 or CD8a T cells was collected and assayed for IFN-γ (R&D Systems, DY485) and IL-10 (R&D Systems, DY417) (CD4 T cells) or just IFN-γ (R&D Systems, DY485) (CD8a T cells) after 3 days of stimulation with αCD3/αCD28, IL-2 and IL-12. Supernatants were assayed according to manufacturer recommendations in duplicate. ELISA plates were analyzed by spectrometer Biotek Synergy H1 plate reader according to the manufacturer’s instructions.

### Generation of bone marrow-derived macrophages and dendritic cells

Bone marrow was isolated from the femurs and tibia of WT and *Aim2^−/−^* mice. Briefly, the bones were cleaned of tissue and flushed with sterile DMEM with 10% FBS and 1% Penicillin/Streptomycin. The red blood cells were lysed with ACK lysis buffer (Gibco, A10492-01). For bone marrow macrophage (BMM) generation, 2 x 10^6^ bone marrow cells were plated on a 10 cm deep dish plate in DMEM with 10% FBS and 1% Penicillin/Streptomycin and 15% CMG14-12-conditioned media for 7 days with additional media being added on day 4. At day 7, BMMs were lifted off and plated on a 24 well plate at 2.5 x 10^5^ cells per well. BMMs were then stimulated for with 100 ng/mL murine IFN-γ (Peprotech, 315-05) for 24 hours. After 24 hours, total RNA was harvested from BMMs, and RT-qPCR was performed as described above.

For the generation of bone marrow derived dendritic cells (BMDDC), 4 x 10^6^ bone marrow cells were plated on a 15 cm plate in RPMI with 10% FBS and 1 % Pentstrep, 50μM β-mercaptoethanol and 20 ng/mL of murine GM-CSF (Peprotech, 315-03) for 10 days with new media being added at day 3, day 6 and day 8. At day 10, BMDDCs were harvested and 2.5 x 10^5^ BMDDCs were plated in a 24 well plate. BMDDCs were then stimulated with 100 ng/mL murine IFN-γ (Peprotech, 315-05) for 24 hours. After 24 hours, total RNA was harvested, and RT-qPCR was performed as described above.

### Statistical Analysis

The GraphPad Prism (Version 10.0.3 for MacOS) was used for analysis. Data are shown as dot plots, where each dot represents an individual mouse/sample and are presented as the mean ± standard error of the mean (SEM).

Data distribution was analyzed using an unpaired student t test for comparisons of 2 means or a one-way ANOVA with Tukey’s post-hoc test for comparisons of >2 means. A *P* value <.05 was considered statistically significant. A ROUT outlier test was used to exclude samples that were deemed outliers.

## Results

### AIM2 Suppresses HNSCC Development

To explore if AIM2 regulates the development of HNSCC, we utilized an established chemical-induced HNSCC mouse model, where Wild type (WT) C57Bl/6J and Aim2-deficient (*Aim2^−/−^*) mice were treated with the chemical carcinogen 4NQO in their drinking water for 20 weeks. WT and *Aim2^−/−^*mice displayed similar weight loss and mortality rates during the 20 weeks of 4NQO treatment (Fig. 1A). After 20 weeks, the tongues of the mice were excised and tumors were enumerated and measured (Figure 1B-1C). While there was no difference observed in the number of macroscopic tumors that developed on the tongues of WT and *Aim2^−/−^*mice, the *Aim2^−/−^* mice displayed significantly larger tumors compared to WT mice (Figure 1B,1C). Tongues from WT and *Aim2^−/−^* mice were fixed in 4% PFA, embedded in paraffin and stained with hematoxylin and eosin for histopathological analysis by a trained investigator blinded to the experimental conditions of the study. In line with larger tumor sizes, *Aim2^−/−^* mice treated with 4NQO were found to have a greater incidence of dysplasia compared to WT mice (Figure 1D-1E), however we did not see any differences in high-grade lesions, severe dysplasia and squamous cell carcinoma. These results suggest that AIM2 plays a restrictive role in oral squamous cell carcinoma initiation but not progression during 4NQO-induced HNSCC.

**Figure 1.**
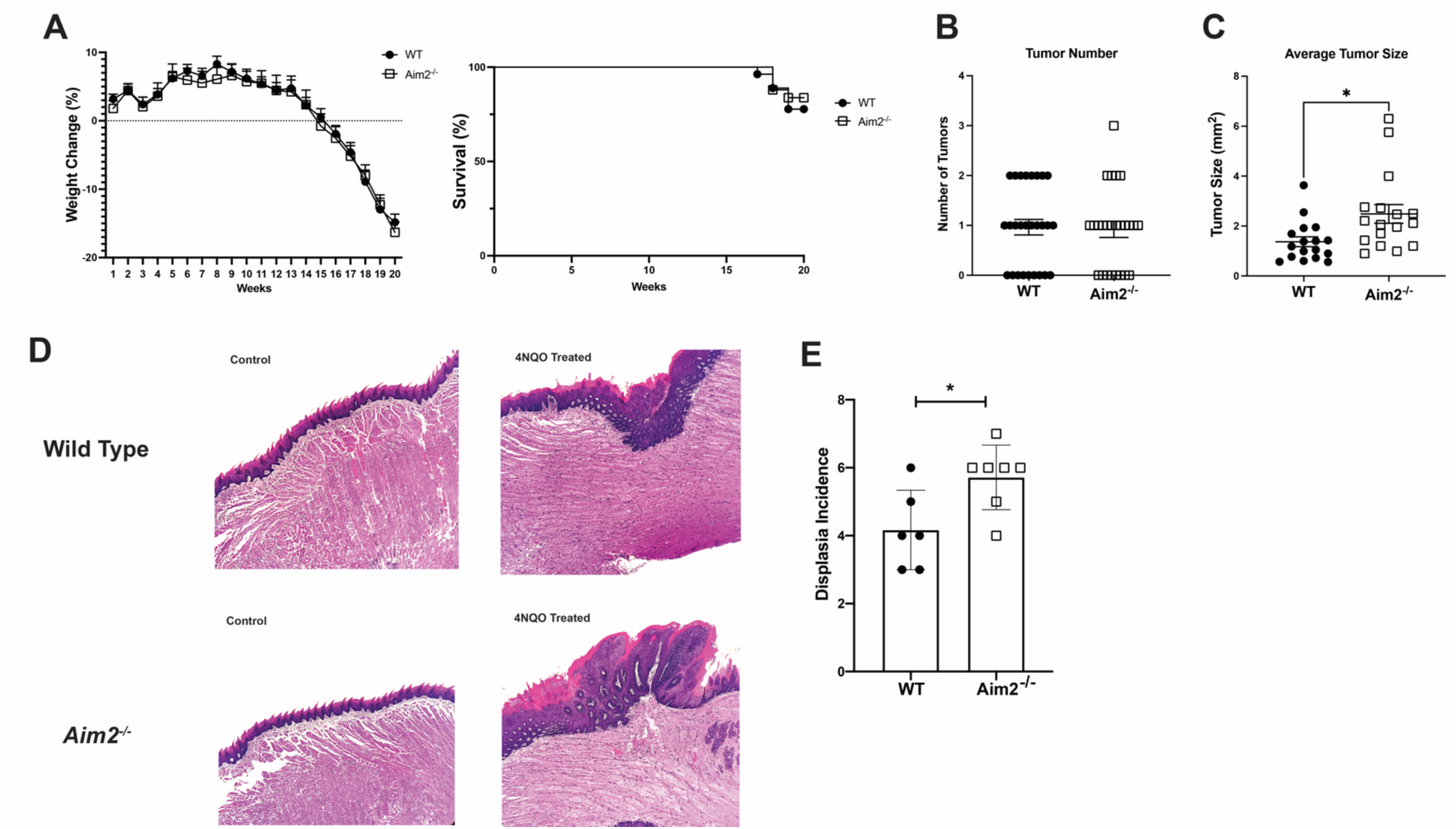
Aim2 restricts tongue tumor size during 4NQO-induced HNSCC. (**A**) Percent weight change (left) and survival (right) of WT (n=21) and *Aim2^−/−^* (n=20) mice treated with 4NQO (left). (**B)** Tumor number enumerated from the tongue of WT (n=21) and *Aim2^−/−^* (n=20) mice (left). (**C**) Macroscopic tumor sizes on tongues of 4NQO-treated WT (n=17) and *Aim2^−/−^* (n=17) mice (right). (**D**) Hematoxylin and eosin staining (H&E) of formalin-fixed and paraffin-embedded WT (n=6) and *Aim2^−/−^* (n=7) mouse tongues. (**E**) Histopathological grading of H&E-stained tongues from 4NQO-treated WT (n=6) and *Aim2^−/−^* mice (n=7). Data are presented as means +/− SEM, *P* values: \**P*<0.05 analyzed by unpaired *t* test unless specified.

### AIM2 limits HNSCC development through inflammasome- and Akt-distinct mechanisms

AIM2 has at least two primary functions, one involving binding to DNA and forming an inflammasome complex with the proteins ASC and Casspase-1 for IL-1β production and the other involving restriction of AKT phosphorylation through interacting with DNA-PKcs. Other inflammasomes promote 4NQO-induced HNSCC^12^, while activating mutations in PI3K/Akt are associated with human HNSCC primarily in HPV^+^ head and neck cancers but is also seen in HPV^−^ cases as well^19^. To assess if AIM2 regulates 4NQO-induced HNSCC through either of these known functions, we excised and sectioned tongues from WT and *Aim2^−/−^*mice treated with 4NQO for 20 weeks and isolated total protein from tongue tissue for western blot analysis of markers associated with inflammasome and Akt activation. To assess inflammasome activation, we western blotted for Caspase-1 to measure levels of active Caspase-1 p20 cleavage product, which is indicative of inflammasome activation. We found similar levels of Caspase-1 cleavage in both WT and *Aim2^−/−^* mice (Figure 2A), indicating overall inflammasome activation was present in the *Aim2^−/−^* mice. We also probed the tongue tissue lysates for phosphorylated AKT by western blot to assess AKT activity. Surprisingly, pAKT levels were reduced in both WT and *Aim2^−/−^*mice given 4NQO, relative to water-treated controls. Moreover, the 4NQO-treated *Aim2^−/−^* mice presenting lower pAKT expression, contrary to the role of AIM2 in limiting Akt activation during other cancers^16^ (Figure 2A). These results indicate that AIM2 limits experimental HNSCC through mechanisms distinct from its known inflammasome-forming and Akt inhibitory functions. Because inflammation is a primary contributor to 4NQO-driven HNSCC^20^, we next tested if AIM2 restricts HNSCC by limiting tissue inflammation. Total RNA was isolated from tongue sections of 4NQO-treated WT and *Aim2^−/−^* mice, converted to cDNA and analyzed for the innate immune cytokines *Il6, Il1b, Tnfa, Il12,* and *Il10* by qPCR (Figure 2C). However, no difference was seen in the expression levels of these cytokines in the WT and *Aim2^−/−^*mouse tongues (Figure 2C), suggesting AIM2 does not impact the expression of common innate inflammatory cytokines during 4NQO-induced tumorigenesis.

**Figure 2.**
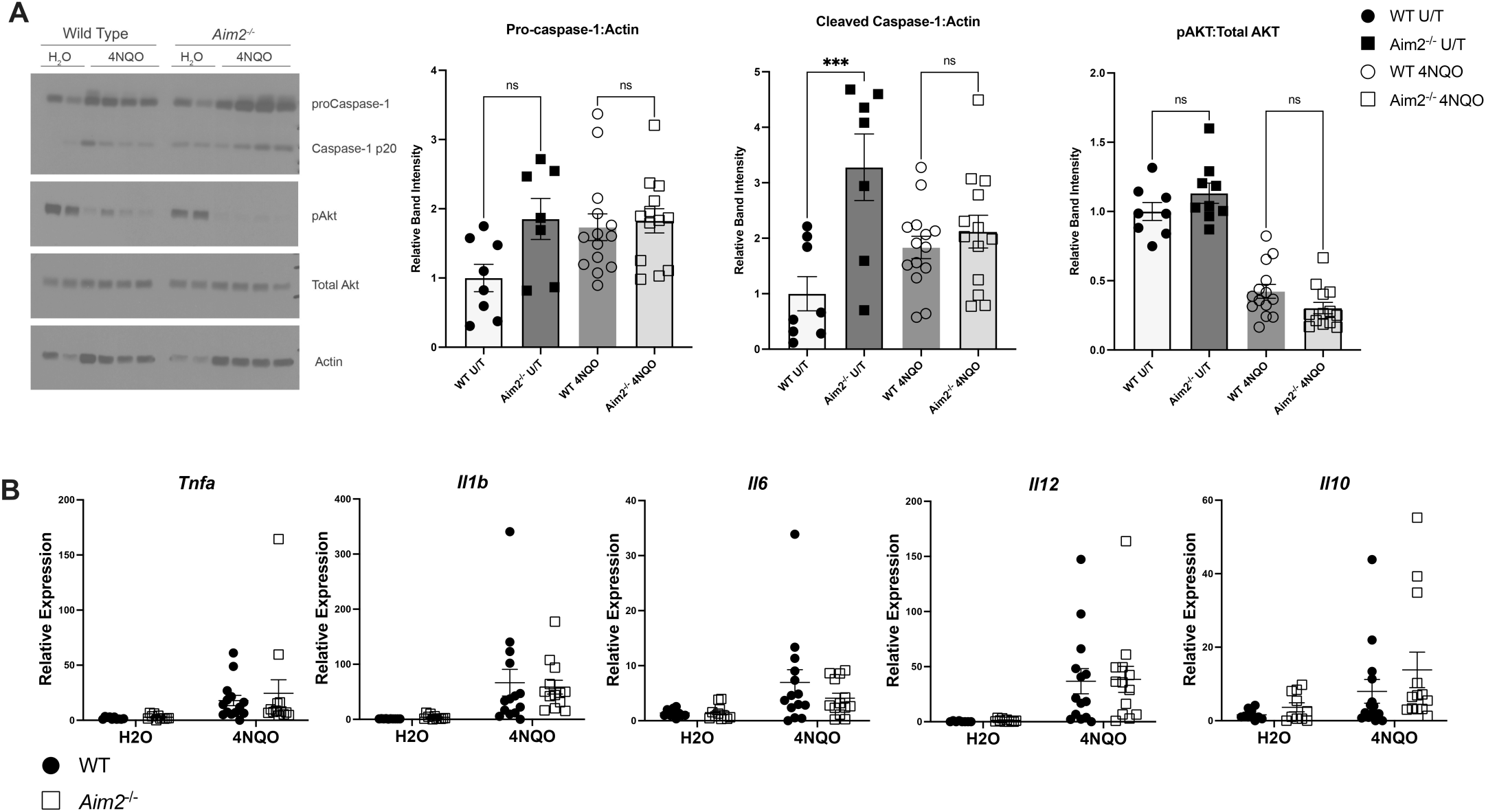
Loss of AIM2 does not result in changes in inflammasome activity, Akt activation or inflammatory cytokine expression during 4NQO exposure. (**A**) (Left) Representative western blot analysis of Caspase-1 cleavage and Akt expression from homogenized whole tongue tissue of WT and *Aim2^−/−^* mice treated with water or 4NQO. Each band represents an individual mouse. (Right) Densitometry analysis of Caspase-1 and cleaved Caspase-1 relative to Actin and phospho-AKT (S473) relative to total AKT from two cohorts of mice. The data was analyzed by one-way ANOVA. P value: \*\*\**P*<0.001. (**B**) RT-qPCR analysis of *Tnfa, Il1b, Il6, Il12,* and *Il10* mRNA expression in tongue tissue from water or 4NQO-treated WT and *Aim2^−/−^* mice n=13-14. Data are presented as means +/− SEM.

### AIM2 Deficiency leads to Elevated IFN-γ Transcriptional Signature during 4NQO Treatment

To gain a better understanding of how AIM2 impacts the transcriptional environment of the tongue in response to prolonged 4NQO treatment, we performed bulk RNA-sequencing on the total RNA isolated from tongue sections of WT and *Aim2^−/−^* mice treated with 4NQO for 20 weeks. We found that *Aim2^−/−^* tongues had 314 unique genes and the WT had 159 (Figure 3A.) Using GO enrichment analysis, we found that genes related to the MHC protein complex, antigen processing and presentation, cell killing and T cell activation were all significantly enriched in the *Aim2^−/−^* tongues (Figure 3A-3C). More specifically, genes encoding H-2A, H-2E, TAP1, Granzyme B and CCR5 were all enriched in the tongues of 4NQO-treated *Aim2^−/−^* mice (Figure 3C). The expression of many of these genes is controlled through several molecular and cellular mechanisms, however the cytokine interferon gamma (IFN-γ) is a potent promoter of many of these genes (e.g., H-2A, H-2E). Conversely, IFN-γ can be a major product of MHC expression and antigen presentation. Therefore, we next measured the expression of IFN-γ in the tongues of 4NQO-treated WT and *Aim2^−/−^* mice and found elevated IFN-γ in *Aim2^−/−^* mice. These findings suggest AIM2 limits production of IFN-γ, which was in line with the increase in antigen processing and presentation genes seen in *Aim2^−/−^*mice (Figure 3B-3C). To confirm this elevated IFN-γ mRNA expression is biologically relevant, we also looked for the tissue expression of *Irf1*, the primary transcription factor activated by IFN-γ signaling. Indeed, *Irf1* expression was significantly elevated in the tongues of 4NQO-treated *Aim2^−/−^* mice (Figure 3D). This suggest that AIM2 limits IFN-γ expression during 4NQO-induced HNSCC.

**Figure 3.**
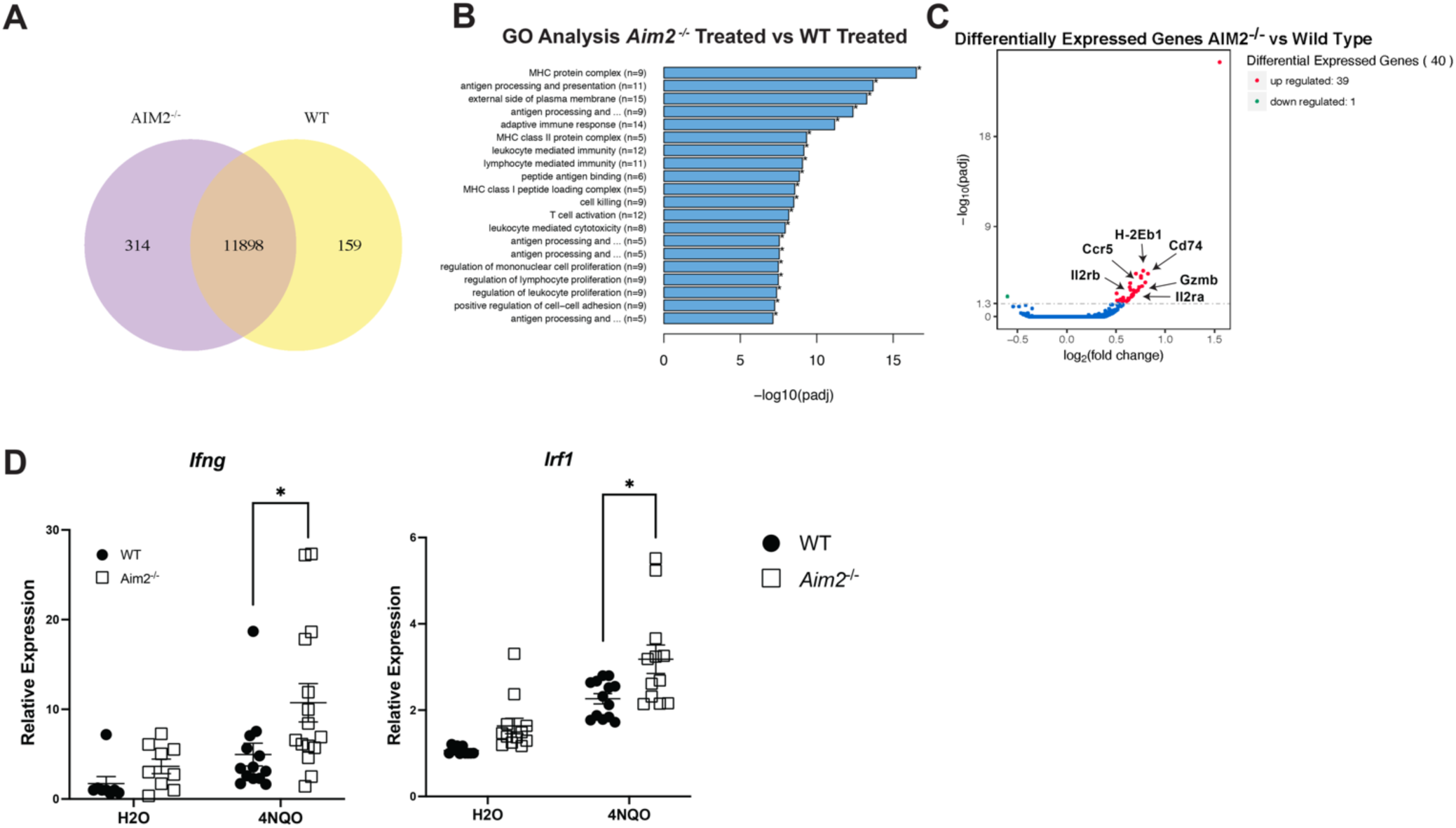
AIM2 Restricts IFNγ During 4NQO-driven HNSCC. Bulk RNA sequencing was performed on mRNA isolated from tongue tissue from 4NQO-treated WT (n=3) and *Aim2^−/−^* (n=3) mice. (**A**) Venn diagram of the number of unique and overlapping genes, **(B)** Gene Ontology (GO) enrichment analysis of transcripts and **(C)** differential gene expression analysis from 4NQO-treated WT vs. *Aim2^−/−^* mice. Arrows indicate genes enriched in *Aim2^−/−^* (n=3) mice treated with 4NQO. (**D**) RT-qPCR analysis of *Ifng* (left) and *Irf1* (right) mRNA expression in tongue tissue from water or 4NQO-treated WT and *Aim2^−/−^* mice n=13-15. Data are presented as means +/− SEM. *P* values: \**P*<.0.05 analyzed by unpaired *t* test unless specified.

### AIM2 deficiency increases macrophage infiltration during 4NQO treatment

We next assessed the immune populations infiltrating the tongue during 4NQO-induced HNSCC through immunohistochemistry staining. Because we detected elevated IFN-γ and genes associated with T cell activation, we first stained 4NQO-treated wild type and *Aim2^−/−^* tongue tissue for CD3e^+^ cells, which is a broad marker for T cells, which are potent IFN-γ producers. We found a trending increase in the number of CD3e^+^ cells infiltrating the epithelium of 4NQO-treated *Aim2^−/−^* tongue tissue, however it did not reach significance (Fig. 4A, 4B). We also found a trending, but not statistically significant, increase in the percentage of CD3e^+^ cells infiltrating the epithelium in 4NQO-treated *Aim2^−/−^* tongue tissue (Fig 4B). These results suggest that loss of AIM2 may lead to a modest increase in infiltrating T cells, but it may not completely explain the elevated IFN-γ observed in the *Aim2^−/−^* mice.

**Figure 4.**
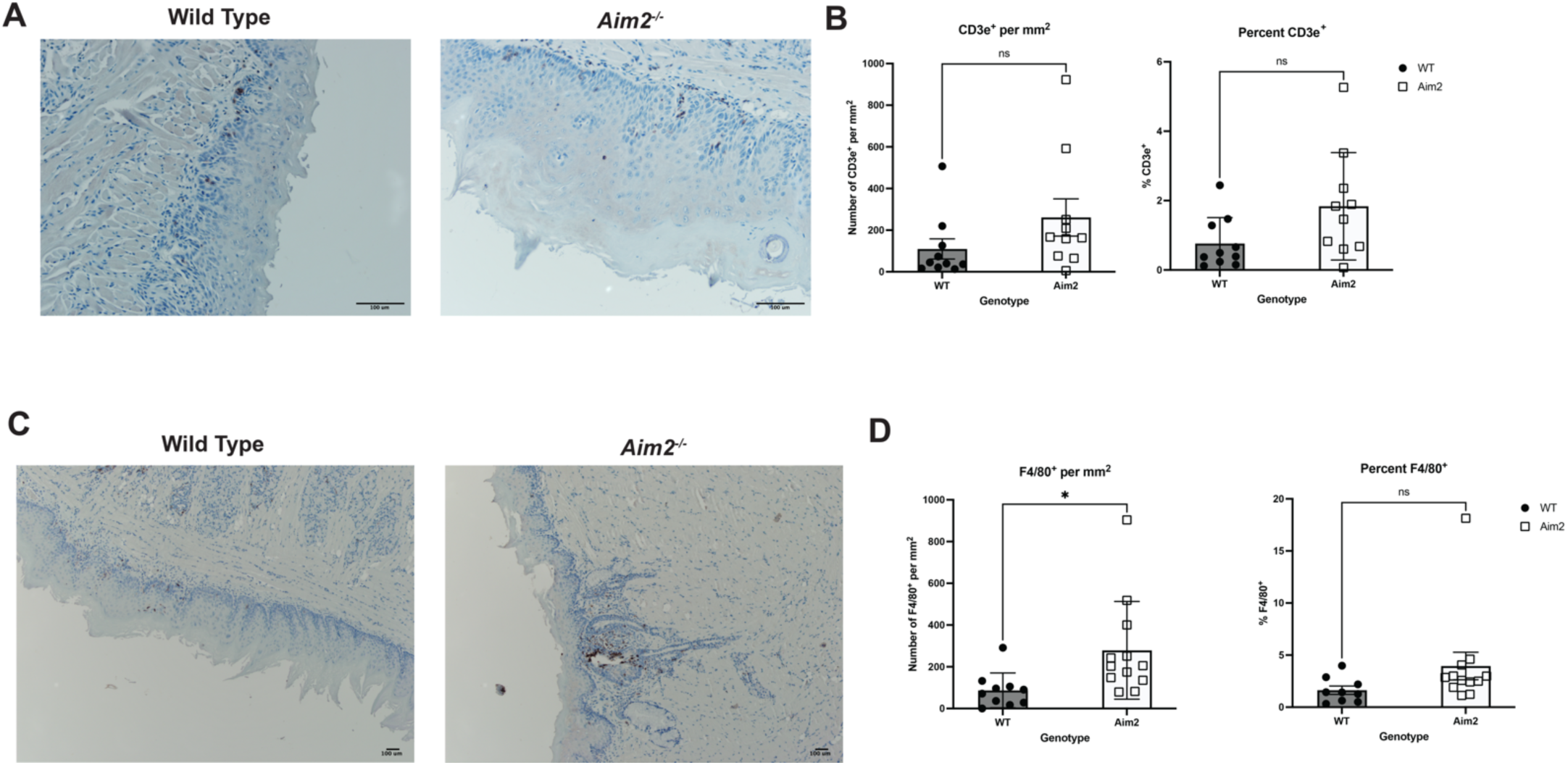
AIM2 deficiency increases tongue macrophage infiltration during 4NQO induced HNSCC. (**A**) Immunohistochemistry analysis of CD3e^+^ cells in tongue epithelium of 4NQO-treated WT and *Aim2^−/−^* mice. (**B**) Quantitative analysis of immunohistochemistry analysis described in (A) of WT (n=10) and Aim2^−/−^ (n=10) tongues. Number of CD3e^+^ cells per mm_2_ (left). Percent of CD3e^+^ cells (right). (**C**) Immunohistochemistry analysis of F4/80^+^ cells in tongue epithelium of 4NQO-treated WT and *Aim2^−/−^* mice. (**D**) Quantitative analysis of immunohistochemistry analysis described in (**D**) of WT (n=10) and Aim2^−/−^ (n=12) tongues. Number of F4/80^+^ cells per mm_2_ (left). Percent of CD3e^+^ cells (right). Data are presented as means +/− SEM. ns, not significant. *P* values: \**P*<0.05 analyzed by unpaired *t* test unless specified.

Next, we stained for the presence of macrophages since we also detected enhanced expression of genes associated with MHC components and antigen processing and presentation in the tongues of 4NQO-treated *Aim2^−/−^*mice. Accordingly, we stained tongue tissue from 4NQO-treated WT and *Aim2^−/−^* mice with F4/80 to measure macrophages (Fig. 4C). We found a significant increase in the number (Fig. 4D left), but not percentage (Fig. 4D right) of F4/80^+^ cells infiltrating the tongue epithelium in *Aim2^−/−^* 4NQO tongue tissue. This suggests that greater numbers of infiltrating macrophages are present in 4NQO-treated *Aim2^−/−^*mice, which may lead to elevated expression of MHC components and antigen processing and presentation genes detected by RNA-seq (Fig. 3C).

### AIM2 expression does not regulate IFN-γ Stimulated Gene Expression in Dendritic Cells or macrophages

The increase in tissue IFN-γ expression in 4NQO-treated *Aim2^−/−^* mice could be explained by two hypotheses; one hypothesis is that AIM2 limits IFN-γ by restricting macrophage and dendritic cell (DC) antigen presentation to T cells. Alternatively, AIM2 may perform an intrinsic function in T cells to limit IFN-γ production. AIM2 is most well described to function in antigen presenting cells like macrophages and dendritic cells^21^. Moreover, we observed an enrichment in genes associated with antigen presentation in 4NQO-treated *Aim2^−/−^* mice (Fig. 3A).

We first asked if AIM2 limits IFN-γ responsive genes in macrophages. These genes include *Irf1*, the primary transcription factor activated by IFN-γ signaling and MHC class II genes*, H-2A* and *H-2E*. We treated bone marrow derived macrophages (BMMs) from wild type and *Aim2^−/−^*mice with IFN-γ *in vitro* for 24 hours and assessed the expression of *Irf1, H-2A* and *H-2E*. We found no difference in the expression of interferon gamma-stimulated genes *Irf1*, *H-2A, H-2E* between WT and *Aim2^−/−^* macrophages (Sup Fig. 1A). This indicates AIM2 does not intrinsically prevent IFN-γ-induced genes in macrophages *in vitro*, which suggests the elevated antigen presentation gene signatures observed in 4NQO-treated *Aim2^−/−^* mice are due to other cellular sources of AIM2. *Aim2^−/−^* dendritic cells, another antigen presenting cell responsive to IFN-γ, also failed to display heightened expression of *Irf1, H-2A* and *H-2E* in response to IFN-γ (Sup Fig. 1B). These findings suggest the overexpression of IFN-γ and interferon-stimulated genes in vivo was not due to an AIM2 intrinsic response by dendritic cells or macrophages (Sup Fig. 1).

### AIM2 does not impact IFN-γ Production in CD4 or CD8 T cells

AIM2 plays a role in regulatory T cell development and function and promotes T helper 17 cell differentiation as well^22–24^. However, the role of AIM2 in CD4 T helper 1 cells (Th1), the predominant IFN-γ producing lineage, is less clear. Specifically, we asked if AIM2 performs an intrinsic function in T cells to limit IFN-γ production. To assess this, we isolated Naïve CD4 T cells from the spleens of WT and *Aim2^−/−^* mice by magnetic activated cell sorting and activated and differentiated them towards the IFN-γ-producing Th1 lineage by treatment with anti-CD3/CD28 antibodies, IL-2, IL-12 and anti-IL-4. At day 4, we isolated total RNA and assessed for *Ifng* expression by RT-qPCR. We found that *Aim2^−/−^* CD4 T cells expressed greater amounts of *Ifng* compared to WT CD4 T cells (Sup Fig. 2A). We also collected cell supernatants from the stimulated CD4 cells and measured IFN-γ protein by ELISA, but found *Aim2^−/−^* CD4 T cells did not secrete more IFN-γ, suggesting AIM2 does not promote IFN-γ production in Th1 cells (Sup Fig. 2A). However, we found that *Aim2^−/−^* CD4 Th1 cells express and produce significantly more IL-10 than WT CD4 Th1 cells, suggesting AIM2 may limit IL-10 production in CD4 Th1 cells (Sup Fig. 2B). There is conflicting data on AIM2’s role in IL-10 production in T cells^22,23^ but our data suggests it restrains its production. We then asked if AIM2 performs an intrinsic function in CD8 T cells to limit IFN-γ production. We isolated Naïve CD8 T cells from the spleens of WT and *Aim2^−/−^* mice by magnetic activated cell sorting and activated them with anti-CD3/CD28 antibodies and IL-12. We then harvested total RNA and supernatants at day 4 and analyzed IFN-γ expression and production. We found no difference in IFN-γ expression in *Aim2^−/−^*CD8 T cells (Sup Fig. 2C). This suggests AIM2 does not play a role in IFN-γ production in either CD4 or CD8 T cells, but may limit IL-10 production in CD4 T cells. Thus, there is no intrinsic defect with the loss of AIM2 causing aberrant IFN-γ production in CD4 and CD8 during 4NQO induced HNSCC.

### Loss of IFN-γ Signaling does not affect tumorigenesis during 4NQO induced Carcinogenesis

Although IFN-γ is often attributed to improved anti-tumor responses, IFN-γ also has tumor-promoting functions^25^. These include selecting for more immunoevasive tumors and contributing towards chronic inflammation and ROS production, which is damaging towards DNA in chronic states and highly associated with cancer development^25^. Moreover, IFN-γ can also promote PD-L1 on tumors^25^ and the loss of T regulatory cells leads to increased IFN-γ and a greater number of 4NQO-driven tumors. ^26^ Importantly, the role of IFN-γ during the early stages of HNSCC development has not been fully explored. To explore the role of IFN-γ in 4NQO-induced HNSCC, we treated WT and IFN-γ-receptor deficient (*Ifngr1^−/−^*) mice to the 20-week 4NQO model (Fig 5). After 20 weeks, the tongues of the mice were excised, and tumors were enumerated and measured as previously described. We found comparable tumor sizes and tumor loads between WT and *Ifngr1^−/−^* mice (Fig. 5A-5B). Moreover, there was also no difference in the number of macroscopic tumors present in the WT and *Ifngr1^−/−^* mice following 4NQO treatment (Fig. 5C). Because loss of IFN-γ receptor failed to result in greater tumor burdens, this data indicates that IFN-γ responses are not solely protective during 4NQO-induced HNSCC development. Moreover, these findings suggest that chronic excessive IFN-γ may worsen 4NQO-induced HNSCC development as observed during Aim2-deficiency.

**Figure 5.**
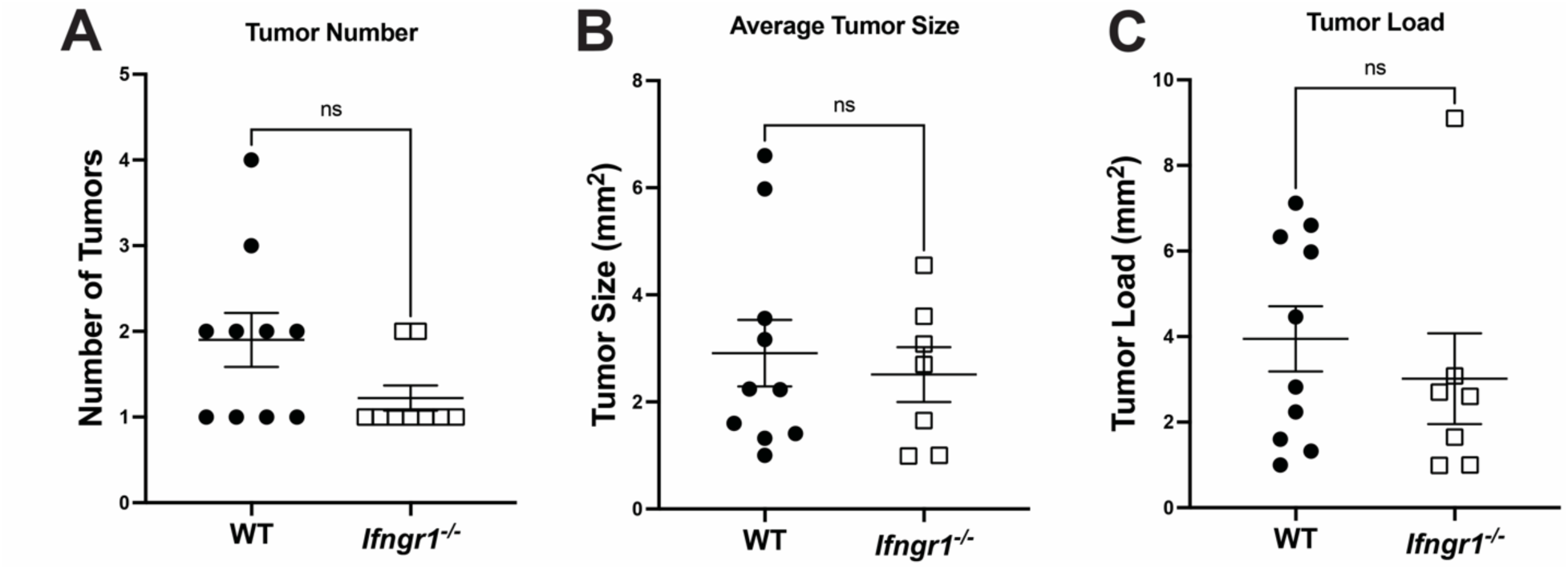
IFNγ is dispensable during 4NQO induced HNSCC. (**A**) Number of macroscopic tumors on tongues of 4NQO-treated WT (n=10) and *Ifngr1^−/−^* (n=7) mice. (**B**) Average tongue tumor sizes in 4NQO-treated WT (n=10) and *Ifngr1^−/−^* (n=7) mice. (**C**) Tumor load on tongues of WT (n=10) and *Ifngr1^−/−^* (n=7). The data were analyzed by unpaired *t* test. Data are presented as means +/− SEM. ns, not significant.

We then wanted to assess the cellular contribution of AIM2 during tumor restriction in vivo. Because we saw elevated IFN-γ in *Aim2^−/−^* mice treated with 4NQO, we assessed the contribution of the adaptive immune compartment during 4NQO-induced HNSCC. We treated *Rag^−/−^* mice, which lack all T and B cells, with 4NQO for 20 weeks. We found the loss of the adaptive immune compartment did not affect the number of tumors, the size of the tumors or the tumor load (Fig. 6A-C). We also treated *Aim2^−/−^/Rag^−/−^*double-deficient mice with 4NQO to determine the contribution of *Aim2^−/−^*in adaptive immune cells during HNSCC development. We found that these mice also showed no difference in tumor numbers, sizes or loads of macroscopic tumors compared to WT mice, failing to recapitulate the elevated tumorigenesis observed in *Aim2^−/−^* mice, which contain a sufficient adaptive immune system (Fig.6A-C). This suggests the adaptive immune system is necessary for the enhanced 4NQO-induced HNSCC in *Aim2^−/−^*mice.

**Figure 6.**
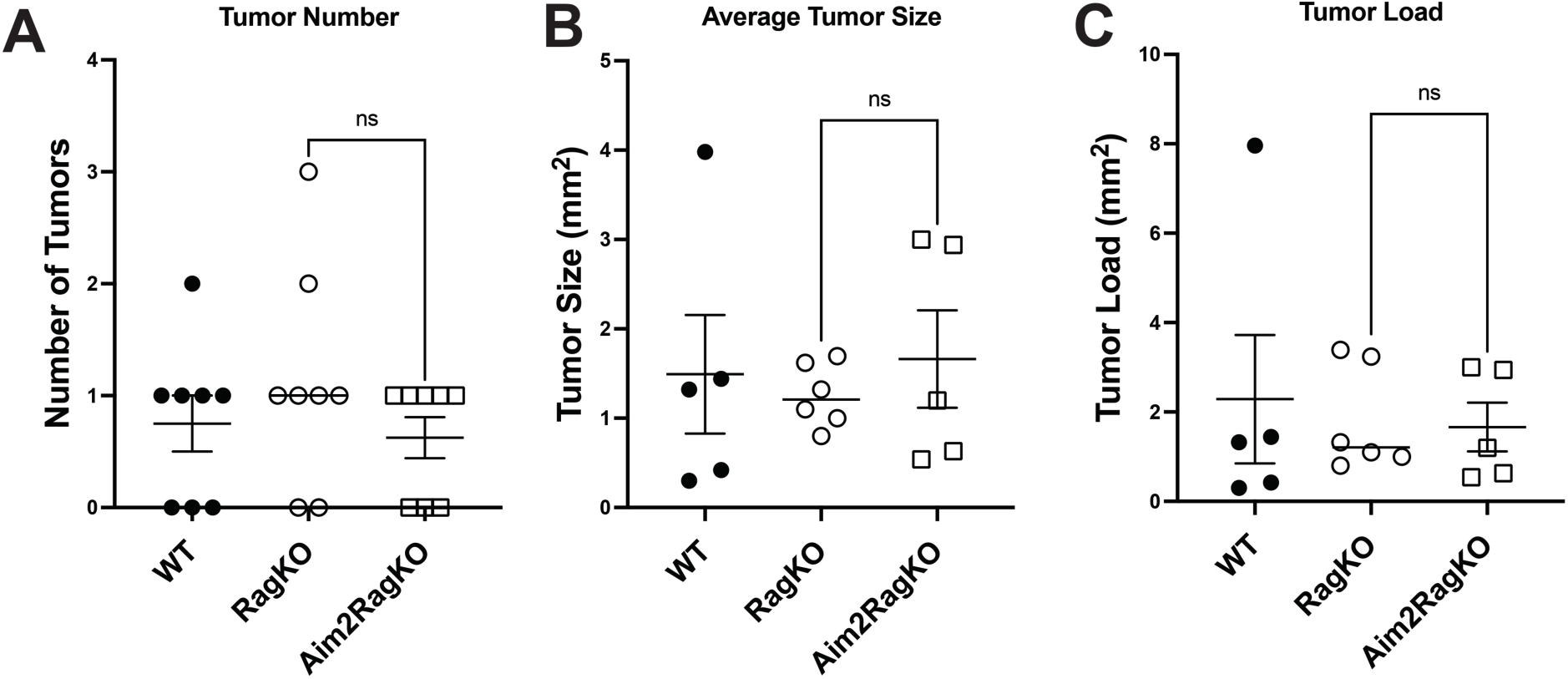
The adaptive immune compartment is necessary for enhanced 4NQO-driven tumor growth during AIM2 deficiency. (**A**) Number of macroscopic tumors on tongues of WT (n=8), *Rag1^−/−^* (n=8) and *Aim2/Rag1^−/−^* (n=8) mice. (**B**) Average tumor sizes on tumors on the tongues of WT (n=5), *RAG1^−/−^* (n=6) and *Aim2/Rag1^−/−^* (n=5) mice. (**C**) Tongue tumor load in 4NQO-treated WT (n=5), *RAG1^−/−^* (n=6) and *Aim2/Rag1^−/−^* (n=5) mice. The data were analyzed by unpaired *t* test. Data are presented as means +/− SEM.

To further assess the specific role of AIM2 in the adaptive immune compartment during HNSCC development, we performed adoptive transfer experiments where we transferred either WT or *Aim2^−/−^* CD4 or CD8 T cells into *Rag^−/−^*recipient mice. We then subjected the mice to 20-week 4NQO treatment. However, no differences in tumor number, size or load were seen between mice that received WT or *Aim2^−/−^* CD4 or CD8 T cells (Fig. 7A-7B). This suggests the enhanced 4NQO-induced HNSCC in *Aim2^−/−^* mice is not driven by AIM2 loss solely by CD4 or CD8 T cells alone and requires the contribution of other or multiple cell types (e.g., epithelium) to fully replicate the enhanced tumorigenesis observed in global AIM2 knockout mice.

**Figure 7.**
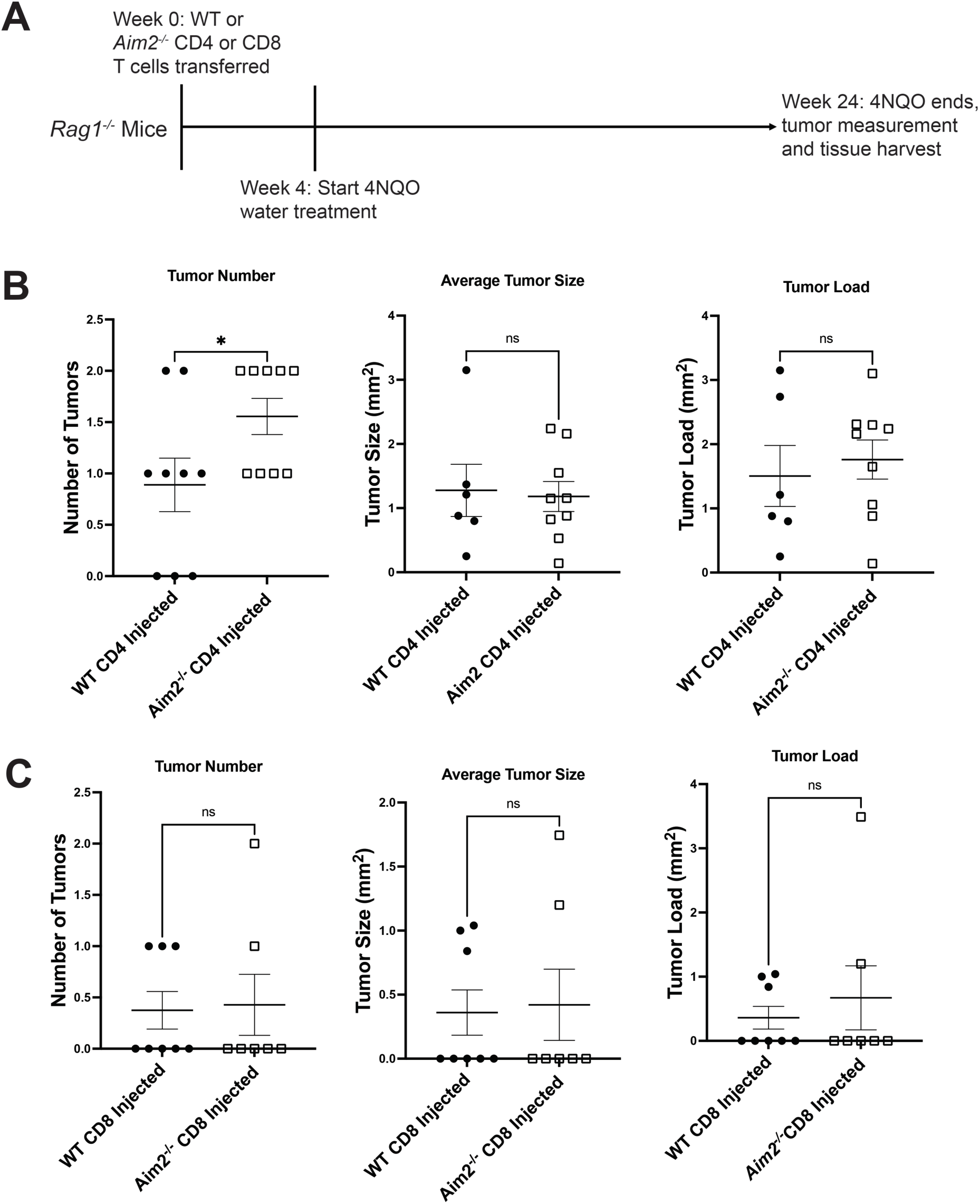
AIM2 Expression in CD4 and CD8 T cells alone are not sufficient for limiting HNSCC. (**A**) Schematic of T cell transfer 4NQO experiments (**B**) Macroscopic tumor number (left), size (middle) and (load) of *Rag^−/−^* mice adoptively transferred WT CD4 T cells (n=9) or *Aim2^−/−^* CD4 T cells (n=9) and given 4NQO. (**C**) Macroscopic tumor number (left), size (middle) and (load) of macroscopic tumors on tongues of *Rag^−/−^* mice adoptively transferred WT CD8 T cells (n=8) or *Aim2^−/−^* CD8 T cells (n=7) and given 4NQO. The data were analyzed by unpaired *t* test. Data are means SEM. ns, not significant.

## Discussion

HNSCC develops in the mouth, nose and throat predominantly due to the carcinogenesis events driven by heavy tobacco and alcohol consumption or HPV infection. Dysregulated immune activation is also major contributor to HNSCC development and progression. However, most of our knowledge base surrounding immune responses during HNSCC is predominantly focused on the tumor microenvironment (TME) of established tumors. This late-stage carcinoma phase of HNSCC is often associated with an immunosuppressive environment. This tumor immune microenvironment is characterized by elevated levels of regulatory T cells (Treg) and tumor-associated macrophages displaying an M2-like phenotype. These anti-inflammatory immune subsets, in combination with tumor-derived immunoregulatory factors (e.g., TGF-β, PGE_2_ and PD-L1) restrict anti-tumor immunity and promote tumor progression^27^. This results in the replacement of effector T cell numbers with exhausted T cell subsets expressing TIM3 and PD-1^28,29^. The blockade of suppressive PD-1 and CTLA-4 proteins on T cells represents an effective immunotherapy for limiting HNSCC development and progression^28,29^. However, while directed immune activation within the TME provides a benefit for a subset of HNSCC patients, chronic or aberrant immune activation is a major contributor during the transition from normal tissue to transformed epithelium. Cigarette smoke, one of the main causes of HNSCC, induces chronic inflammation on mucosal surfaces^30^. It does this by primarily damaging the epithelial cells resulting in inflammatory cytokine release, upregulation of certain PRRs and activation of NF-κB that promotes chronic immune cell recruitment to the area and inflammation^31–33^. In particular, IL-1β can promote malignant transformation and aggressiveness in HNSCC, further showing the importance of inflammatory cytokines in early tumor initiation and progression^14^. However, there is still much to be explored during these early events of tumor initiation.

The 4NQO model of HNSCC models the development of tobacco or chemical induced head and neck cancer. ^34^ This model shows similar histological and genetic alterations that are seen with tobacco induced HNSCC^34^. This model also induces chronic inflammation, which is an important driving factor for HNSCC. ^34^ Several PRRs promote inflammation and many have been linked to the initiation and development of cancer. This includes the inflammasome-forming protein NLRP3, which responds to a broad range of microbial components, like the pore-forming toxin nigericin, and damage-associated host components like extracellular ATP^12,13^. In the 4NQO model of HNSCC, loss of NLRP3 delayed tumor growth and relieved the immunosuppressive tumor state^12^. In this model, NLRP3 promotes tumor growth through the chronic release of IL-1β. IL-1β is elevated in both human tumors and oral tissue from animals treated with 4NQO, and elevated IL-1β is associated with worse tumor outcomes^14^. Several other inflammasome-forming proteins exist and have been linked to other cancers, but their role in HNSCC development as not been explored.

AIM2 represents another inflammasome-forming PRR that responds to cytosolic DNA for the release of the inflammatory cytokines IL-1β and IL-18^15^. AIM2 also functions as a negative regulator of AKT phosphorylation, which results in suppression of colon tumorigenesis^16^. Conversely, AIM2 can drive inflammation during carcinogenic liver injury^35^. However, AIM2’s role in head and neck cancer is less clear as well as if either the inflammasome-promoting and/or Akt suppressive functions of AIM2 are involved. Recent work found that in established HNSCC tumors and cell lines, AIM2 is upregulated and represents a poor prognostic marker in tumors^17^. In this study, AIM2 expression was linked to promoting tumor radio resistance, migration and PD-L1 and was associated with altered STAT1 and NF-κB activity^17^. Moreover, low expression of AIM2 along with high expression of STAT3 was associated with poor prognosis of hypopharyngeal squamous cell carcinoma^36^, which is related, but anatomically distinct from oral cancer. Importantly, both studies focus on the expression of AIM2 on late stage HSNCC without assessing its contribution on the immune system. Here, we found that during the initiation of HNSCC, the loss of AIM2 lead to larger tumors, an elevated IFN-γ transcriptional signature and greater infiltration of macrophages to the epithelium. These combined data suggest AIM2 may play opposing roles during HNSCC initiation versus progression, a process characterized by distinct immune signatures and activation states.

Our work sought to determine if AIM2 plays a role in HNSCC carcinogenesis. Utilizing a 20-week 4NQO model, we found that *Aim2^−/−^* mice grew larger tumors compared to similarly treated WT mice. These mice also had a greater incidence of dysplasia suggesting that during early HNSCC transformation/development, AIM2 plays a suppressive role. To our surprise, we did not see differences in the expression of many innate inflammatory cytokines in the tongue of *Aim2^−/−^*mice including TNF, IL-6 and IL-1β. No measurable loss of caspase-1 cleavage was observed in *Aim2^−/−^* mice, suggesting global inflammasome function was intact in these mice, likely through the presence of other inflammasome sensors. Because other inflammasomes, like the NLRP3 inflammasome, promotes tumorigenesis in response to 4NQO, we concluded that the inflammasome function of AIM2 was not involved in this process. We next hypothesized that AIM2 performed a suppressive role during HNSCC through its ability to suppress AKT as described in colorectal cancer^16^. The PI3K/AKT pathway is one of the most frequently mutated pathways in HNSCC, and tumors that harbor these mutations have a higher rate of mutations in cancer-related genes^6^. However, upon assessing Akt activation, we found that *Aim2^−/−^* mice did not display increased phosphorylation of Akt in tongue tissue compared to similarly treated WT mice, suggesting AIM2 may not suppress HNSCC through limiting Akt.

While AIM2 failed to alter the expression of many innate inflammatory cytokines during 4NQO exposure, we did observe a heightened IFN-γ transcriptional signature in the tongues of *Aim2^−/−^* mice. This includes elevated expression of the cytokine IFN-γ and IFN-γ-stimulated genes such as MHC class II proteins, TAP1 and granzyme B. This suggests a role for AIM2 directly or indirectly in suppressing either IFN-γ responses or production during 4NQO induced HNSCC. Interestingly, *Aim2^−/−^* mice also present elevated IFN-γ during models of gastric cancer through an inflammasome-distinct mechanism^37^. Also we observed increased IFN-γ in the *Aim2^−/−^* water treated control mice compared to wild type, suggesting that there may be elevated levels of IFN-γ at baseline or it may be due to the aging of the mice since these mice were age matched with the 4NQO treated mice. IFN-γ is a type 2 Interferon that is often considered protective during cancer and a marker of beneficial anti-tumor immunity, however it also has pro-tumoral effects depending on the context^25^. For example, IFN-γ can induce the expression of the immunosuppressive protein PD-L1^38^, select for more invasive and metastatic tumors^39^ and induce the generation of reactive oxygen species, which can directly damage DNA leading to mutations during chronic inflammation^40^. In the context of early HNSCC development, the role of IFN-γ is not as clear. Deletion of Tregs leads to aberrant effector T cell activation and IFN-γ production, which promotes tumorigenesis during 4NQO-induced HNSCC ^26^. This indicates that unrestrained IFN-γ is detrimental and not protective during the early stages of HNSCC development. In line with this finding, *Ifngr1^−/−^* mice given 4NQO displayed similar tumor burdens as WT mice, suggesting IFN-γ is not solely protective during early tissue transformation. However, further work is needed to clarify the multifaceted role of IFN-γ during HNSCC development versus progression.

IFN-γ production is tightly controlled and produced by specific cells; these include activated CD4^+^ and CD8^+^ T cells as well as Natural killer cells, ILC1s and γδ T cells. The majority of research focusing on AIM2 is related to its role in innate immune cells and its inflammasome function. As we did not detect major differences in innate cytokines present in *Aim2^−/−^* tissue, but instead found robust IFN-γ and related gene expression profiles, we evaluated the role of AIM2 in the context of the adaptive immune system. The role of AIM2 in lymphocytes is only beginning to be explored. AIM2 is thought to play a role in CD8^+^ T cells during gastric cancer^37^. AIM2 also contributes to Treg stability and development^22,23^, but opposing roles for AIM2 have been identified during this process. One group found that AIM2 suppresses Treg cell differentiation through modulating T cell metabolism^23^, while another group revealed that AIM2 functions in regulatory T cells to restrain autoimmunity by reducing Akt-mTOR signaling and enhancing the stability of Treg populations^22^. In the context of HNSCC, Tregs restrain effector T cell responses and prevent exacerbation of HNSCC^26^. We did not find altered IFN-γ production when *Aim2^−/−^* CD4^+^ or CD8^+^ T cells were activated ex vivo, indicating no intrinsic role for AIM2 in suppressing IFNγ production from these cells. However, *Aim2^−/−^* CD4^+^ T cells did show elevated levels of IL-10 suggesting a role for AIM2 in limiting IL-10 production during Th1 skewing. This finding is in line with that of Lozano et. al. that found the loss of AIM2 leads to increased IL-10 in Tregs^23^, suggesting a suppressive role for AIM2 during IL-10 production of IL-10. We did not find elevated numbers of T cells in the tongues of treated *Aim2^−/−^* mice however, the composition of the infiltrating T cells was not determined. We next directly asked if the loss of AIM2 in the adaptive immune compartment was driving the increased tumor sizes in *Aim*2^−/−^ mice during 4NQO induced HNSCC. *Aim*2^−/−^ mice crossed with lymphocyte-deficient RAG1^−/−^ mice displayed no enhancement of tumor burdens, suggesting that lymphocytes are required for the increased tumorigenesis observed in 4NQO-treated *Aim*2^−/−^ mice. However, adoptive transfer of WT or *Aim2^−/−^* CD4^+^ T cells into RAG1-/- recipient mice lead to no observable differences during 4NQO-induced tumorigenesis, suggesting loss of AIM2 expression in CD4 T cells alone was not sufficient to promote this enhanced tumor burden. We also conducted a similar experiment using adoptive transfer of WT or *Aim2^−/−^* CD8^+^ T cells into RAG1^−/−^ mice and found similar results. These experiments suggest that AIM2 expression in CD4^+^ or CD8^+^ T cell populations alone are not sufficient to control HNSCC. Similar findings were observed for the involvement of AIM2 during experimental gastric cancer study where deletion of AIM2 in several individual cell types was insufficient in recapitulating the increased pathology observed in the global AIM2 deficient animals^37^. Because AIM2 is expressed by several immune and epithelial types that are involved in tumorigenesis, these results suggest that other (or multiple) cell types may contribute to HNSCC restriction in the globally deficient *Aim2^−/−^*mice.

Macrophages are a major cell type that respond to IFN-γ and they can display pro- or anti-tumor activities depending on their environmental exposure. Generally, M1 macrophages mediate anti-tumor effects through direct killing, with one mechanism being through production of reactive oxygen and nitrogen species^41^. However, M2 macrophages are typically considered cancer promotors primarily through their production of various cytokines that suppress anti-tumor immunity and stimulate tumor cell growth and survival^41^. IFN-γ promotes M1-like responses by directly inducing anti-microbial ROS/RNS, increasing antigen presentation to T cells by inducing expression of Major histocompatibility (MHC) genes and recruiting innate and adaptive immune cells through chemokine production^42^. RNAseq revealed that *Aim2^−/−^* mice treated with 4NQO displayed elevated expression of genes linked to antigen processing and presentation (i.e., *Tap1*, *H-2eb1*, *H-2ab1,* and *B2m)* and chemotaxis (i.e., *Ccr5*), which are associated with M1 macrophage-like activity. In addition, we observed an increase in macrophage infiltration into the tongue epithelium of the 4NQO-treated *Aim2^−/−^*mice. However, we observed that *Aim2^−/−^* macrophages stimulated with IFN-γ in vitro did not have altered IFN-γ-induced gene signatures, including *Irf1, H-2eb1* and *H-2ab1*. This suggests that the heightened IFN-γ signatures in the 4NQO-treated *Aim2^−/−^* mice may be due to elevated tissue infiltration of macrophages versus an intrinsic role AIM2 in these cells during the response to IFN-γ. Dendritic cells, another important antigen presenting cell type that responds to IFN-γ, also displayed no differences in these IFN-γ-stimulated genes when AIM2 was absent. Therefore, exact cellular and molecular mechanisms by which AIM2 limits IFN-γ and related gene signatures remains to be clarified during HNSCC development. Future work will ascertain if macrophages in conjunction with other IFN-γ producers and responders are responsible for the increases in tumor sizes in 4NQO-treated *Aim2^−/−^* mice. Moreover, it will be important to understand the impact of AIM2-regulated IFN-γ expression during HNSCC progression and within the tumor-immune microenvironment of established tumors.

## Acknowledgments

The authors would like to thank the University of Arizona Cancer Center’s Experimental Mouse Shared Resource (EMSR) and Tissue Acquisition and Cellular/Molecular Analysis Shared Resource (TACMASR) for their technical assistance. Both shared resources were supported by the Cancer Center Support Grant **P30 CA023074**.

## Footnotes

This work was supported by Institutional Research Grant number IRG-16-124-37 from the American Cancer Society (to J.E.W.), National Institutes of Health, National Cancer Institute Grants K22CA212030 (to J.E.W.), T32CA009213 (to D.M.R), and R01CA274857 (to C.C.) and the National Institutes of Health, National Institute of Dental Craniofacial Research Grant 5F31DE032263 (to D.M.R).

## Author Contribution Statement

D.M.R and J.E.W designed the experiments and wrote the paper; D.M.R and S.L.T performed the majority of the experiments; V.E-R. performed the IHC staining and assessments, with supervision from C.C.; C.C. performed the histopathological assessments, discussed the data and provided critical review of the manuscript.

