## Supplemental Figures 1-2 for "Impact of AIM2 on HNSCC Development"

**Supplementary Figure 1.**

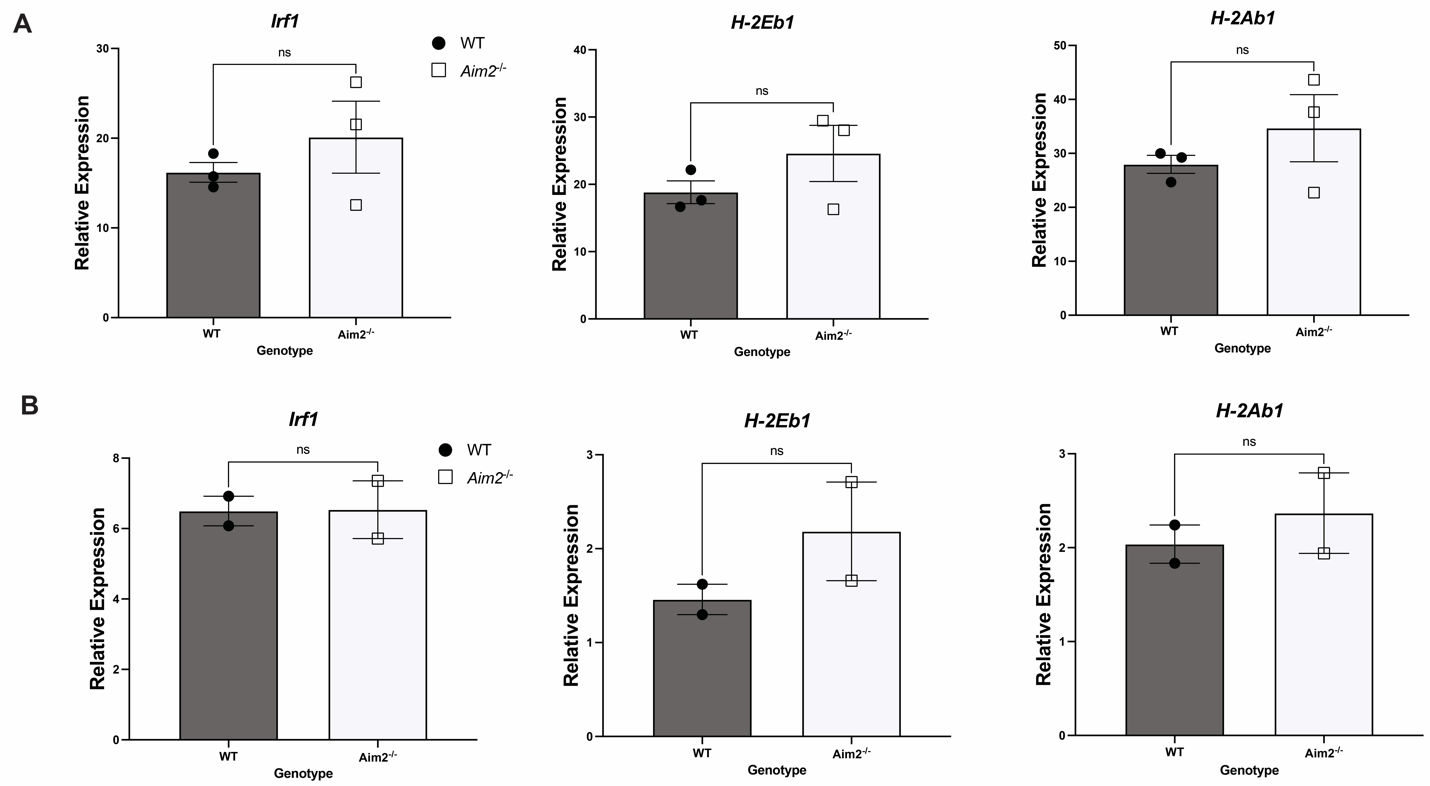

**Supplementary Figure 1. AIM2 does not regulate *Irf1* or MHCII expression in macrophages or Dendritic cells in vitro.** (**A**) RT-qPCR analysis of *Irf1, H-2Eb1* and *H-2Ab1* mRNA expression from WT (n=3) and *Aim2^-/-^* (n=3) bone marrow-derived macrophages stimulated with IFN-𝛾 for 24 hours. (**B**) RT-qPCR analysis of *Irf1, H-2Eb1* and *H-2Ab1* mRNA expression from WT (n=3) and *Aim2^-/-^* (n=3) bone marrow-derived dendritic cells stimulated with IFN-𝛾 for 24 hours. Data are presented as means +/- SEM. ns, not significant by unpaired *t* test.

**Supplementary Figure 2.**

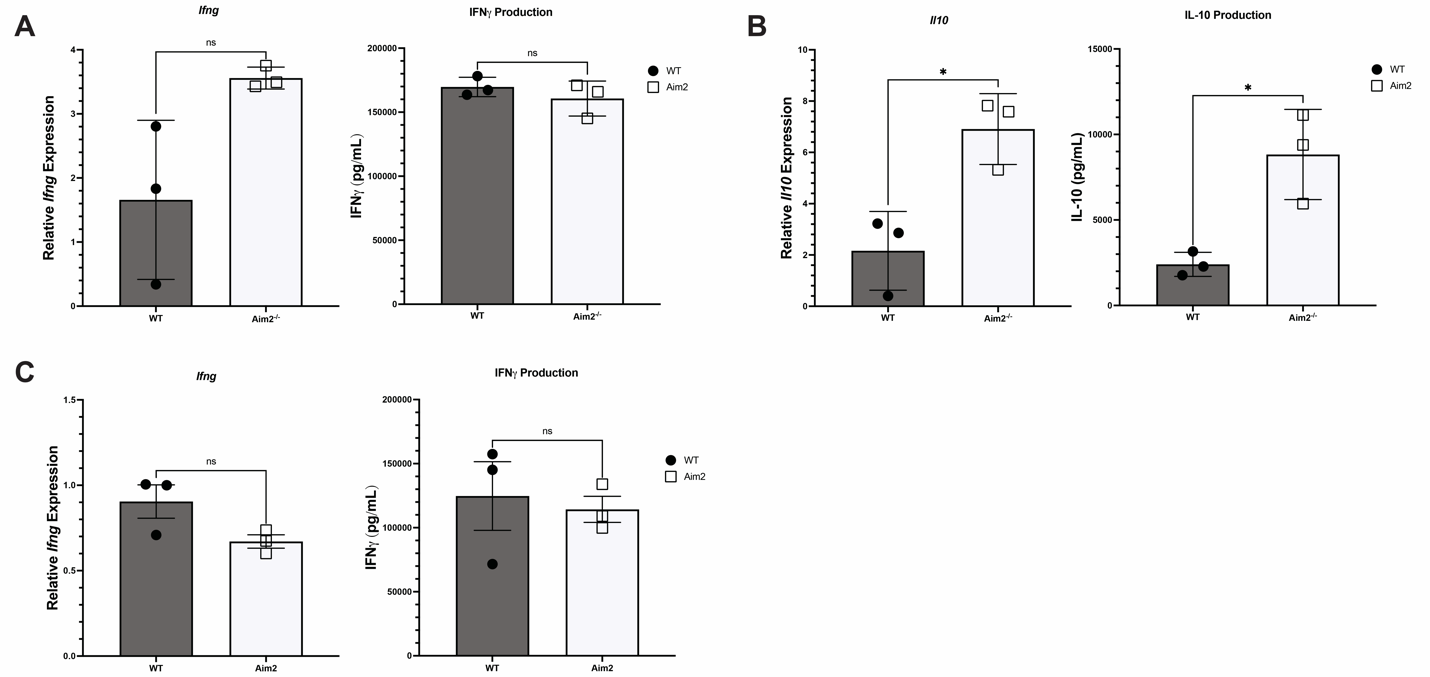

**Supplementary Figure 2. No intrinsic role for AIM2 in restricting IFN𝛾 in CD4+ or CD8+ T cells in vitro.** (**A**) RT-qPCR analysis of *Ifng* mRNA expression (left) and IFN-𝛾 levels in culture supernatants (right) from WT (n=3) and *Aim2^-/-^* (n=3) naïve splenic CD4^+^ T cells activated ex vivo with αCD3/CD28, IL-2 and IL-12 for 3 days. (**B**) RT-qPCR analysis of *Il10* mRNA expression (left) and IL-10 levels in supernatants (right) from WT (n=3) and *Aim2^-/-^* (n=3) naïve splenic CD4^+^ T cells activated ex vivo with αCD3/CD28, IL-2 and IL-12 for 3 days. (**C**) RT-qPCR analysis of *Ifng* mRNA expression (left) and IFN-𝛾 levels in culture supernatants (right) from WT (n=3) and *Aim2^-/-^* (n=3) naïve splenic CD8^+^ T cells activated ex vivo with CD3/CD28, IL-2 and IL-12 for 3 days. Data are presented as means +/-SEM, *P* values: **P*<0.05, ***P*<0.01 analyzed by unpaired *t* test.
